# A possible stimulus to induce SPAWNING in Ezo abalone *Haliotis discus hannai* during stormy weather: Fenton reaction induces spawning behavior

**DOI:** 10.1101/2020.10.30.356469

**Authors:** Yukio Matsumoto, Kousuke Yatsuya

## Abstract

Synchronous spawning is an important behavior to increase fertilization success of invertebrates with external fertilization. Previous work has shown that it is possible to induce spawning behavior using free radicals in tank experiments, but the stimulus for spawning in the wild is not fully understood. Ezo abalone *Haliotis discus hannai* mainly spawn during stormy weather. Rainwater contains H_2_O_2_ and iron (II) ions (Fe^2+^). We propose that during stormy weather water layers in the ocean are mixed and the surface layer containing H_2_O_2_ and Fe^2+^ interacts with the ocean bottom; this leads to conditions suitable for the Fenton reaction to occur. Hydroxyl radicals (·OH) are generated during the oxidization of Fe^2+^ by H_2_O_2_ and we hypothesized these induce spawning behavior of abalone in the wild. This study observed that *H. discus hannai* released eggs after salinity decreased due to the rainfall during stormy conditions. In addition, our tank experiment demonstrated that ·OH generated by the Fenton reaction induced synchronous spawning behavior between the sexes. This study provides a new hypothesis about control of synchronous spawning in *H. discus hannai* and the results could be applicable to other invertebrates.

## 1. Introduction

Synchronous spawning is an important behavior to increase fertilization success of marine invertebrates with external fertilization. Although the circalunar rhythm [1], stormy weather [2–4], high increased phytoplankton concentration [5,6], and tidal change [7] could synchronize gamete release between opposite sexes, the primary stimulus for releasing gametes is not known for many species. Under laboratory conditions, free radicals that react with organic matter are able to induce spawning behavior in many marine invertebrate [8–14]. Morse et al., 1977 [15] demonstrated that *Haliotis rufescen* spawned when a high concentration of hydrogen peroxide (5mM H_2_O_2_) was added to the tank; in this example, H_2_O_2_ served as a donor of highly reactive species such as hydroperoxyl radicals and peroxyl radicals. These radicals increased prostaglandin synthetase, which subsequently induced egg maturation and spawning behavior.

The spawning of *Haliotis* species in Japan is observed during stormy weather [2–4]. Rainwater and sea surface water contain low concentrations of H_2_O_2_ (rainwater, [16,17]; sea surface, [18]) and iron (II) ions (rainwater, [19,20]; sea surface, [21]), which are sources of hydroxyl radicals (·OH) through the Fenton reaction. In this reaction, iron (II) ions (Fe^2+^) are oxidized to iron (III) ions by H_2_O_2_, and ·OH and hydroxide ions are generated [22–24]. The stormy weather could mix layers of water, allowing the surface layer containing H_2_O_2_ and Fe^2+^ to reach the sea bottom. We hypothesized that the Fenton reaction occurs on the sea bottom, and the ·OH generated by the Fenton reaction could induce the spawning behavior of abalone in the wild. First, this study investigated whether Ezo abalone *H. discus hannai* released eggs in the wild after a decrease in salinity caused by rainwater. Second, we tested the effect of the Fenton reaction on the spawning behavior of *H. discus hannai* in tank experiments.

## 2. Methods

### Study species

Ezo abalone *Haliotis discus hannai* is a marine gastropod mollusk in the family Haliotidae, which is distributed in the waters off Japan and eastern Asia. The species mainly inhabits rocky shores. The spawning season at our study site in the Iwate Prefecture, Japan (39° 36’ N, 142° 2’ E), is from August to late October, when water temperatures are around 20–21°C.

### Field observations of egg release

To investigate the effect of salinity changes on the release of eggs, we attached data loggers (length: 7cm, φ: 1.5cm; weight: 23g; Biologging solutions Inc.) to female abalone to record egg releasing behavior. Females swing their shells during egg release, and therefore egg release could be characterized by changes in acceleration data [25]. Test females were captured at the study site and were kept in a tank for approximately 1 wk. The gonad index was determined based on previously developed methods [26], and abalone with mature gonads were used for the study. A plastic frame for mounting the data logger was attached to the abalone shells using epoxy resin. The data logger was mounted onto the plastic frame using cable ties one day before releasing. Two females fitted with data loggers were released at 09:00 AM on 12th August 2016 (ID1 and ID2) and 3 females (ID3, ID4, and ID5) were released at 09:00 AM on 3rd September 2016 (start logging at 9:00 AM on 6th September). The females were released by SCUBA divers at 10 m depth. ID5 was not recaptured. The salinity and water temperature per 10 min at 10 m depth were recorded using A7CT-USB (JFE Advantec, Japan). Atmospheric pressure and rainfall in Miyako, about 10 km northwest of the study site, was provided by the Japan Meteorological Agency. The acceleration data from abalone behavior were analyzed using Ethographer (https://sites.google.com/site/ethographer/docs). Accelerations with specific frequencies (x-axis, 4.9-11 s; y-axis, 1-12.3 s; z-axis, 1.5-11.5 s) and amplitudes (x-axis, 0.076-0.74 m/s^2^; y-axis, 0.073-0.713 m/s^2^; z-axis, 0.12-0.45 m/s^2^) were extracted as putative egg releasing behavior. Matsumoto et al., 2018 [25] described the detailed procedure for acceleration data analysis. The details of the abalones used in the study are presented in ESM. 1.

### Effect of the Fenton reaction on gamete release

We tested the effect of hydroxyl radicals (·OH) on induction of spawning behavior in *H. discus hannai* from 2nd February to 8th March, 2020. The details of the abalone used in this experiment are presented in ESM. 2. The abalone were reared in a stock tank (2500 L) at 20 °C prior to the experiments. Abalone are able to release gametes after an effective accumulated temperature (EAT) of 1000 degree-days [26]. For the experiment, we selected abalone with mature gonads from tanks that had reached 1500-2000 degree days. We filled the experimental tanks with water from a stock tank to minimize the effect of change in water quality on the release of gametes. Spawning experiments were conducted after sunset when *H. discus hannai* are most active. Each experimental abalone was moved to a separate experimental tank (20 L) at 13:00, and the abalone were acclimated to the experimental tanks until 17:30-18:00. A heater and aeration were used to keep water temperature at 20 °C. The concentrations of H_2_O_2_ and Fe^2+^ in the tank were determined based on the concentrations observed in the wild. We first added H_2_O_2_ into the experimental tank followed by Fe^2+^. The final concentration of H_2_O_2_ (40 μM; FUJIFILM Wako Pure Chemical Corporation, Japan) was within the range of rain water (8.4-82μM) [27] because the concedntration of H_2_O_2_ in the seawater during stormy condition is not clear. There is a possibility that the Fe^2+^ concentration of rainwater decreases after mixing with seawater. However, Fe^2+^ could also be added from ground water and rivers [28]. Since the Fe^2+^ concentration of seawater during stormy conditions cannot be calculated exactly, the concentration of Fe^2+^ (55 nM) in this experiment was within the range of rainwater during stormy conditions (12-102 nM) [29]. Fe^2+^ exists as an organoiron complex in sea water [30,31] and rainwater [31]. In this experiment, readily available iron citrate was used as a spawning inducer. Stock iron citrate (Fe^2+^-citrate) was made by dissolving 3 g citric acid (KENEI Pharmaceutical Co., Ltd., Japan) and 160 mg iron (II) chloride (NACALAI TESQUE, INC., Japan) with 1.5L ultrapure water, and 75 μl stock iron citrate was added into the experimental tank (the Fenton group). The Fe^2+^-citrate solution was prepared on the day of experimentation to prevent the oxidation of the Fe^2+^. To test the effect of a low concentration of H_2_O_2_ in our experiment, we set up tanks with 40 μM H_2_O_2_ (the H_2_O_2_ group). In addition, we had a control tank without any reagents to account for any impact of handling stress (the control group). The experimental tanks were placed in the dark. We confirmed that the experimentation tanks produced a higher concentration of ·OH (ESM. 3). We observed the tanks every 30-60 minutes and recorded if gametes were observed. The period from the addition of chemicals to the observation of spawning is regarded as the time before spawning. The proportion of abalone releasing gametes was defined as egg release rate and sperm release rate. Egg release rate and sperm release rate were analyzed with a Generalized liner model (GLM) with binomial distribution followed by Shaffer post-hoc test using “multcomp” package for R v3.6.3.

## 3. Results

### Field observation of egg release

Putative egg release behavior was not detected from ID1 and ID2, however, putative egg release behavior was detected from ID3 and ID4. We could also determine whether the females released their eggs by observing the gonads visible through their shells. ID1 and ID2 did not release eggs and ID3 and ID4 released their eggs before being recaptured. The putative egg release behavior was observed at 11:27 on 6th September from ID3 and at 19:17 on 9th September from ID4. The sea conditions during the study period are presented in ESM.4, Fig. 1a, and Fig. 1b. The study site was calm from August 12th to 15th (ESM. 4_1), and then a low-pressure system came to the study site on September 6th and 8th (ESM. 4_2). The salinity decreased about 3 hours before spawning of ID3 (Fig. 1b and 1c). The spawning of ID4 was also observed during low salinity (Fig. 1b and 1d).

**Fig.1.**
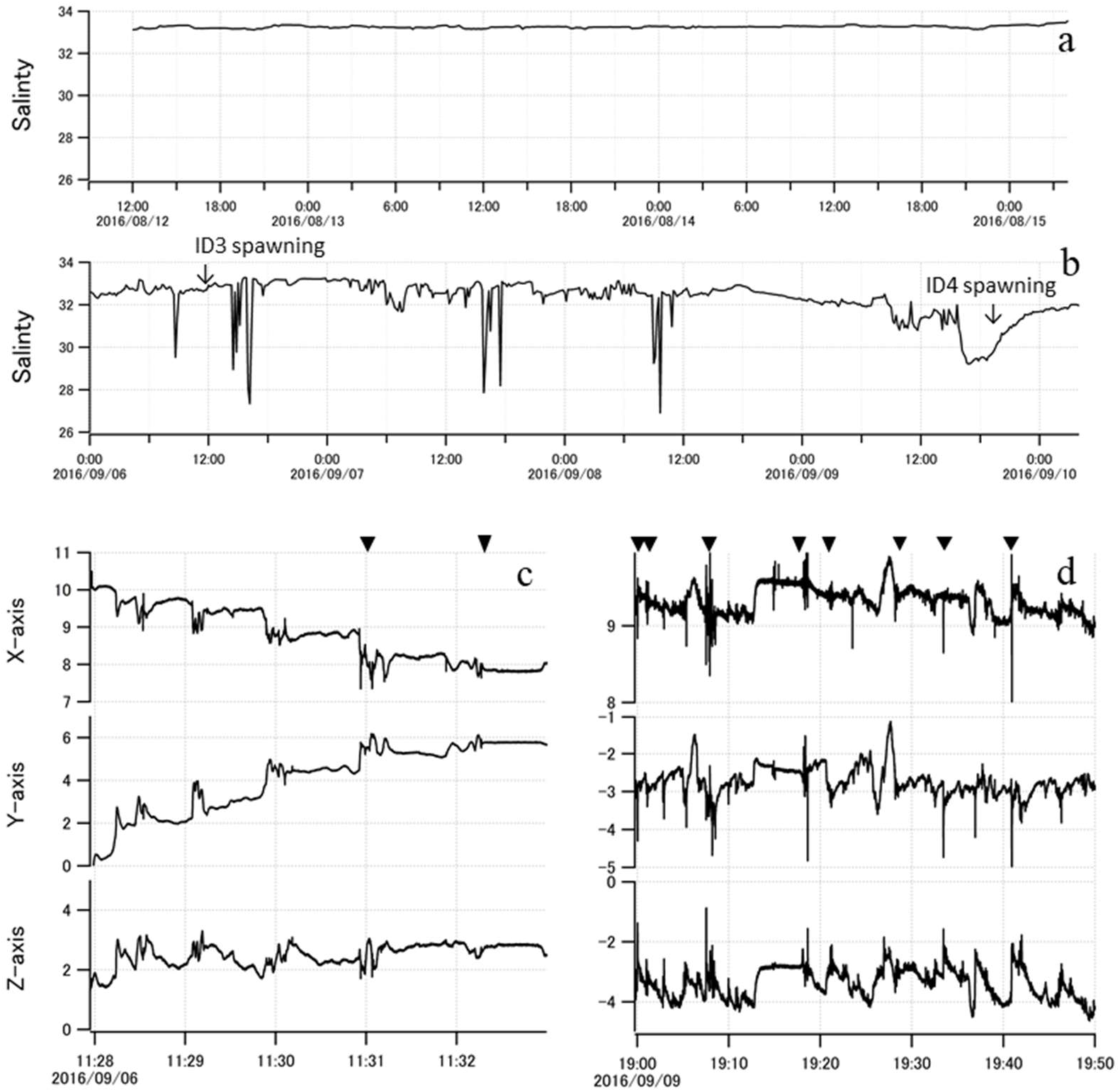
The relationship between salinity and egg releasing behavior. Time series of salinity at 10 m depth when (a) ID1 and ID2 were tested and (b) when ID3 and ID4 were tested. Time series acceleration data of (c) ID3 and (d) ID4 when putative egg release behaviors were observed. Arrow heads indicate the putative egg release behavior.

### Effect of the Fenton reaction on gamete release

The egg release rate of the Fenton group was higher than the H_2_O_2_ group and control group (GLM with Shaffer test, Fenton vs. H_2_O_2_, *z* = 3.34, *p* = 0.002; Fenton vs. control, *z* = 2.86, *p* = 0.004; H_2_O_2_ vs. control, z = 0.66, *p* = 0.50; Fig. 2a). There was no difference in sperm release rate among groups (Fenton vs. H_2_O_2_, *z* = 0.54, *p* = 0.59; Fenton vs. control, *z* = 1.85, *p* = 0.20; H_2_O_2_ vs. control, z = 1.24, *p* = 0.20; Fig. 2b), however, the sperm release rate in the Fenton group was higher than that of other groups. The test abalone in the Fenton group released their gametes and therefore their gonads were smaller than at the start of the experiment. The amount of gametes released in the control and H_2_O_2_ groups was very small. The time before gamete release for females and males in the Fenton group ranged from 60-180 min, however, the time before egg release in the H_2_O_2_ group and the control group was more widely dispersed (Fig. 2c).

**Fig.2.**
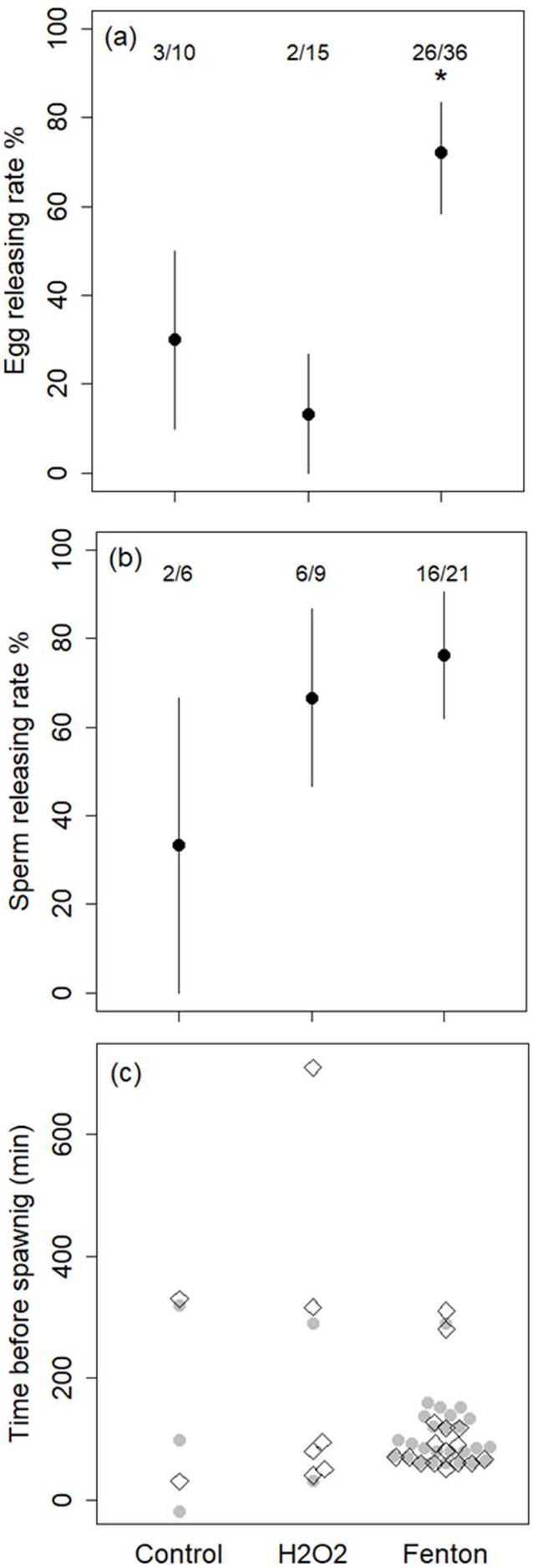
The comparison of (a) egg release rate, (b) sperm release rate, and (c) time before spawning among the Control, H_2_O_2_, and Fenton group. Error bars indicate the prediction interval based on the binomial distribution. ● indicates female abalone and ◇ indicates male abalone. The numbers above the plot indicate the number of abalone spawning and the number of trials in each experiment (No. spawning abalone / No. of trials).

## Discussion

This study indicated a possible primary stimulus to induce spawning of *H. discus hannai.* In the wild, putative egg releasing behavior was observed after salinity decreased. The decreasing salinity could indicate that the rainwater containing H_2_O_2_ and Fe^2+^, which are sources of ·OH through the Fenton reaction, reached the sea bottom. The tank experiments in this study showed that synchronous spawning occurred after the addition of H_2_O_2_ and Fe^2+^ that is consistent with possible concentrations in the wild during stormy conditions. These results strongly suggest that rainwater containing H_2_O_2_ and Fe^2+^ can reach the sea bottom and induce synchronous spawning through the Fenton reaction. Changes in water temperature in the wild (ESM. 4_2. e) may also induce spawning as suggested by research of *H. discus discus* [32]. However, the induction of spawning by temperature change was not synchronous among the sexes under laboratory conditions [33]. There is the possibility that handling stress prior to and at release may induce spawning in *H. discus hannai.* However, since putative spawning was observed 3-6 days after release of the test abalone, there should be a low possibility that handling stress influenced the results. The logger started logging 3 days after release, and therefore the effect of handling stress on spawning before logging was initiated cannot be assessed in this study.

It is likely that·OH generated by the Fenton reaction induced spawning in the tank experiments. Time before spawning (60-180 min) was consistent with the time required for final oocyte maturation; in female *Haliotis* species final oocyte maturation occurs after addition of the spawning inducer [34], followed by egg release [33] after 60-90 minutes at the earliest [35]. Although male gonads have sperm that are able to fertilize eggs [36], males release sperm after 60-90 minutes to synchronize with female egg release [35]. The observed egg release within 60 min in the H_2_O_2_ and control treatments may have been induced by other stimulations (i.e., handling stress). Measurement of ·OH under stormy conditions in the future will help to support the validity of our tank experiment.

When a weaker low-pressure system passed on 6th September, putative spawning of ID3 was observed, however, ID4 did not show any spawning behavior. In *H. discus hannai*, the extent of spawning is correlated with wave height [2]. Spawning of only ID3 during the weaker low-pressure system may have demonstrated small-scale spawning. As the amount of rainfall before spawning of ID3 was smaller than that of ID4 (ESM. 4), a lower concentration of ·OH may explain small-scale spawning. The behavior of ·OH in the sea may also have influenced the effect on ID4. Even though ID4 was exposed to low salinity several times before 9th September, the salinity decrease was experienced in the daytime. Spawning rate of this species under culture was 40% lower during the day relative to the night [35]. The circadian rhythm of abalones may explain the absence of ID4 spawning before 9th September. On the other hand, there could be an additional stimulus for gamete release during a large-scale storm, other than the amount of ·OH. To demonstrate our hypothesis, temporal and spatial variation of the Fenton reaction should be investigated. In addition, more observation of spawning behavior of abalone in the wild is needed.

This study presents a new hypothesis that connects previous observations that free-radicals induce spawning (e.g., [15,35]) with a source of free-radicals in the wild. Our hypothesis may identify the primary stimulus for spawning in other invertebrates. The spawning of the sea urchin *Strongylocentrotus nudus,* the gastropod, *tegla spp.,* and the mytilid, *Septifer virgatus* was indeed in sync with *H. discus hanani* [2]. As the surface water contains H_2_O_2_ [18] and Fe^2+^ [21], tidal-dependent spawning of other *Haliotis* species [7] may occur when the surface water containing H_2_O_2_ and Fe^2+^ reaches the sea bottom. Not only the Fenton reaction but also production of free radicals by phytoplankton [37] and solar radiation [38] could generate a spawning cue. Algae induced spawning [5,6] may be explained by the free radicals derived from phytoplankton. The understanding of the sources of free radicals in the wild would help to further understanding of the spawning biology of invertebrates in the wild.

## Supporting information

ESM

## Acknowledgements

Messrs. Yuichi Kohsaka and Nao Kimura of Omoe Fisheries Cooperative Association helped us carry out the field investigation. We thank Fukumi Tashiro for her help in rearing the experimental animals. We also thank Hideki Takami for his comments on the manuscript.

