## Supplementary material for "A possible stimulus to induce SPAWNING in Ezo abalone *Haliotis discus hannai* during stormy weather: Fenton reaction induces spawning behavior": ESM: ESM.1.docx

ESM.1. Information of the abalones attached with data-logger. Raw data have been deposited in Mendeley Data at <https://data.mendeley.com/datasets/2xtcpkgfcd/draft?a=9a4b38c5-4d9a-495b-8ac9-4a6ae949716c>.

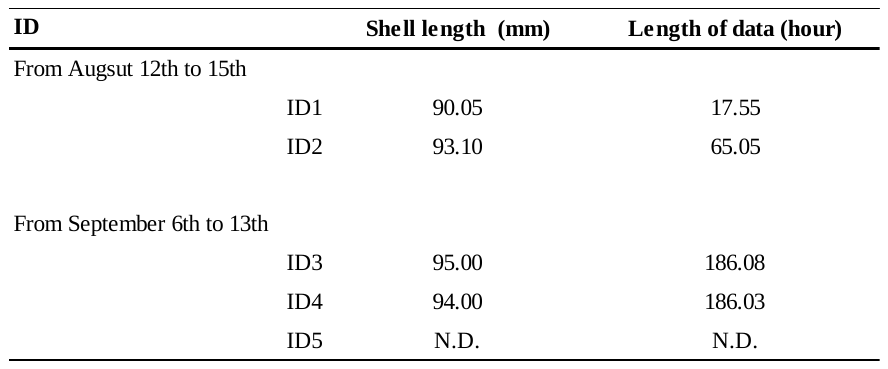
