## Supplementary material for "A possible stimulus to induce SPAWNING in Ezo abalone *Haliotis discus hannai* during stormy weather: Fenton reaction induces spawning behavior": ESM: ESM.2.docx

ESM.2. Information of the abalones in the tank experiments. Cultured abalone (> 4 years old) from the Japan Fisheries Research and Education Agency (Miyako laboratory) and abalone captured off Iwate Prefecture in November 2019 were used in this experiment.

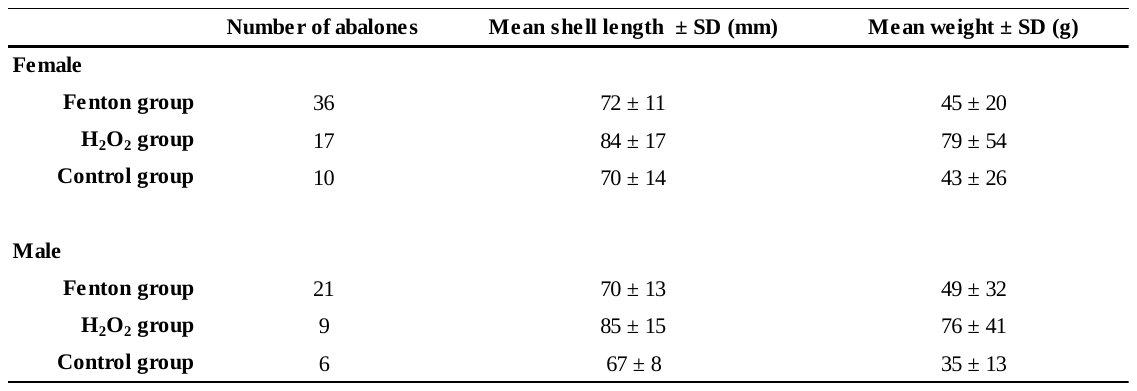
