## Supplementary material for "A possible stimulus to induce SPAWNING in Ezo abalone *Haliotis discus hannai* during stormy weather: Fenton reaction induces spawning behavior": ESM: ESM.3.docx

ESM. 3. Observation of Hydroxy radicals

**Methods**

To confirm that Hydroxyl radicals (^･^OH) were generated in the Fenton group, behavior of ^･^OH was observed using a fluorescence probe on June 12, 2020. This experiment was conducted in a 20 ml plastic tube, which is 1/1000 scale of the tanks used for the spawning experiments.^･^OH converts terephthalic acid (TA) to 2-hydroxyterephthalic acid (HTA) that can be detected by measurement of fluorescence ( e.g., Saran and Summer, 1999). HTA emits light at 425nm when it is irradiated by 310nm light (e.g., Kanazawa et al., 2011). The concentration of ^・^OH was estimated using the fluorescence strength of HTA, because the ^・^OH trap rate in TA in our experimental condition was not clear. The quantification was carried out by following the steps described below. The reagents used in the experiments were prepared within 24 hours of measurement. For this measurement, the H_2_O_2_ and Fe^2+^-citrate were diluted to 1/1000 with ultrapure water. As TA does not dissolve in acidic/neutral liquid, we dissolved TA (FUJIFILM Wako Pure Chemical Corporation, Japan) in a 1M NaOH (FUJIFILM Wako Pure Chemical Corporation, Japan). The concentration of TA solution was 200 mM. H_2_O_2_ was added into the sample tube filled with 20 ml of filtered seawater to make a concentration of 40 μM. We added the TA solution (final concentration 0.2mM) and Fe^2+^*-*citrate (final concentration 55 nM) into the sample tube in that order. The addition of TA solution did not increase pH of the sample solution. The sample solution was measured using a spectrofluorometer (FP-8500, JASCO Corporation, Japan). The production of ^・^OH was observed immediately after addition of Fe^2+^*-*citrate, at 10 minutes, 30 minutes, and 1 hour.

**Results**

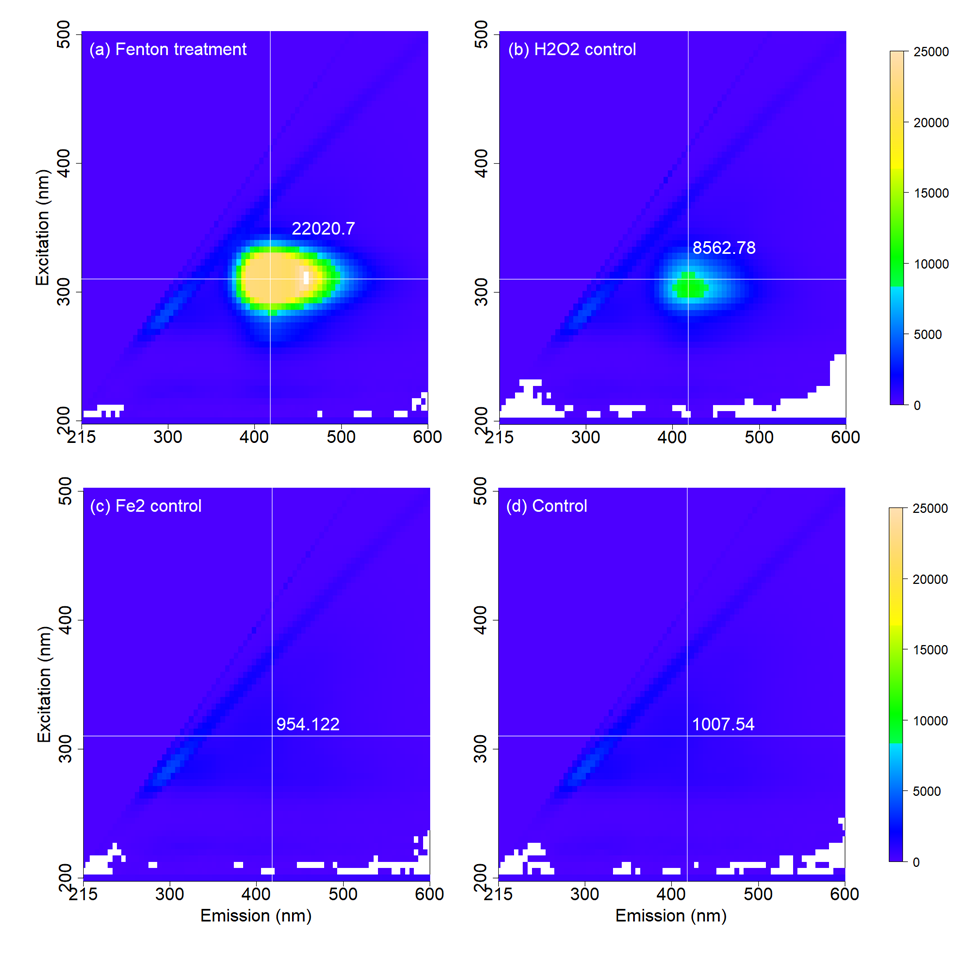
^･^OH was observed immediately after Fe^2+^*-*citrate addition (Figure. a). In the H_2_O_2_ control (Figure. b), Fe^2+^ control (Figure. c), and the control group (Figure. d), ・OH was not detected.
