## Supplementary material for "A possible stimulus to induce SPAWNING in Ezo abalone *Haliotis discus hannai* during stormy weather: Fenton reaction induces spawning behavior": ESM: ESM.4.pptx

### Slide 1
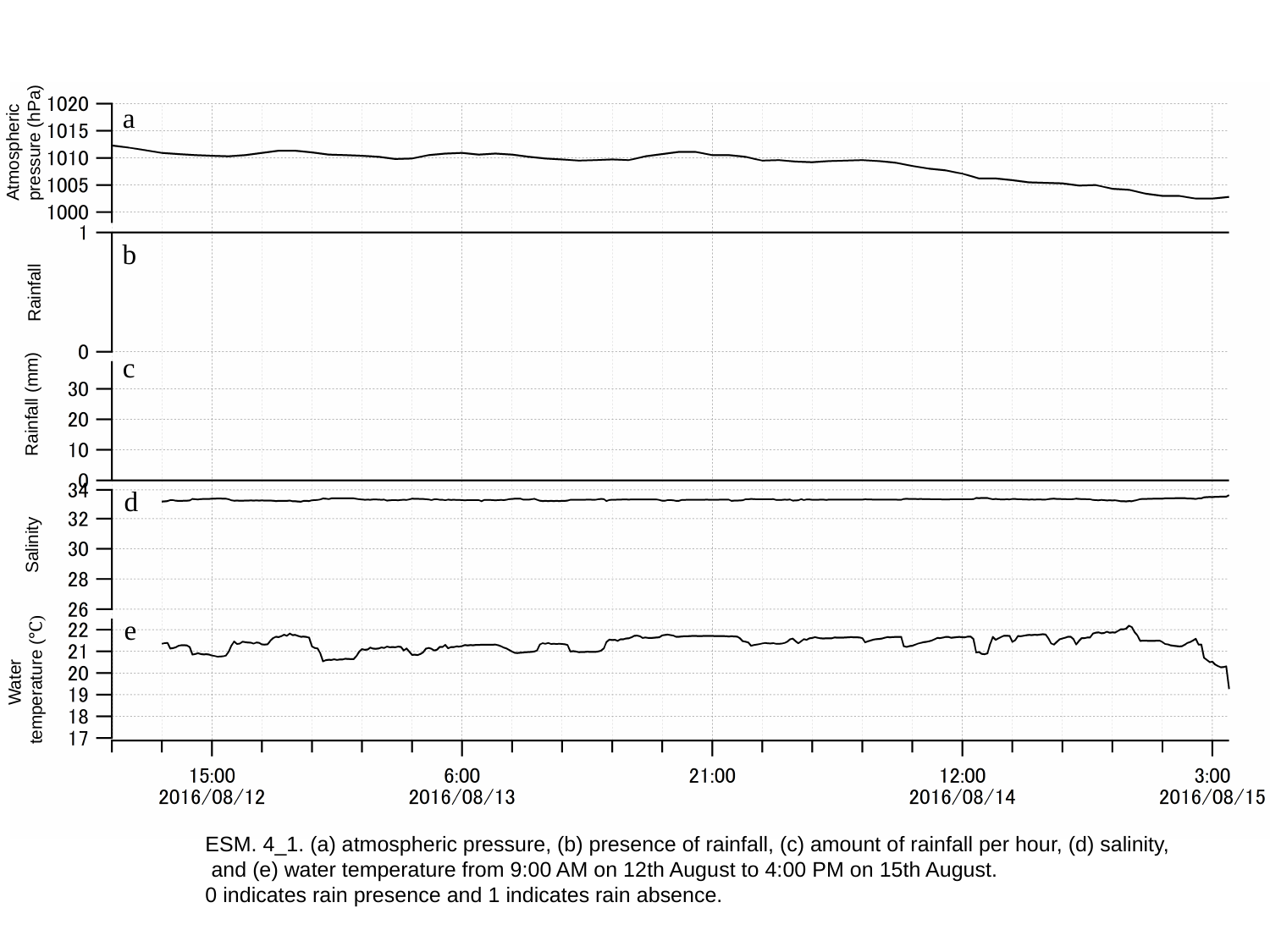

a
Atmospheric
pressure (hPa)
b
Rainfall
c
Rainfall (mm)
d
Salinity
e
Water
temperature (℃)
ESM. 4_1. (a) atmospheric pressure, (b) presence of rainfall, (c) amount of rainfall per hour, (d) salinity,
 and (e) water temperature from 9:00 AM on 12th August to 4:00 PM on 15th August.
0 indicates rain presence and 1 indicates rain absence.

### Slide 2
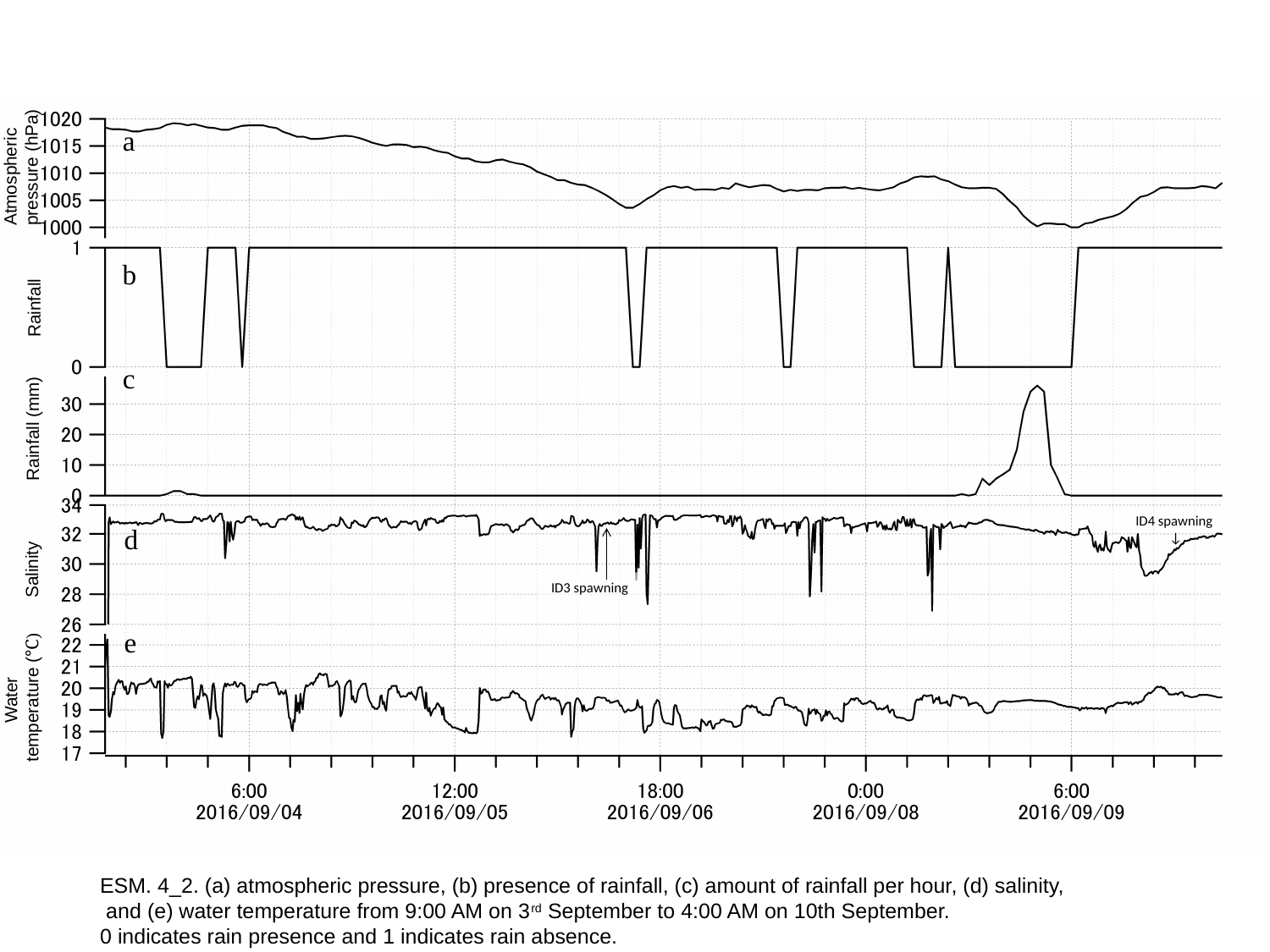

a
Atmospheric
pressure (hPa)
b
Rainfall
c
Rainfall (mm)
ID4 spawning
↓
d
Salinity
ID3 spawning
e
Water
temperature (℃)
ESM. 4_2. (a) atmospheric pressure, (b) presence of rainfall, (c) amount of rainfall per hour, (d) salinity,
 and (e) water temperature from 9:00 AM on 3^rd^ September to 4:00 AM on 10th September.
0 indicates rain presence and 1 indicates rain absence.
